# Targeting dCas9-SunTag to a susceptibility gene promoter is sufficient for CRISPR interference

**DOI:** 10.1101/2025.03.11.642632

**Authors:** Zuh-Jyh Daniel Lin, Gabriela L. Hernandez, Myia K. Stanton, Xingguo Zheng, Kerrigan B. Gilbert, Suhua Feng, Basudev Ghoshal, Jason Gardiner, Ming Wang, Steven E. Jacobsen, James C. Carrington, Rebecca S. Bart

**Affiliations:** Donald Danforth Plant Science Center, Saint Louis, MO 63132, USA; Department of Molecular, Cell and Developmental Biology, University of California, Los Angeles, CA 90095, USA; Eli & Edythe Broad Center of Regenerative Medicine & Stem Cell Research, University of California, Los Angeles, CA 90095, USA; Sidney Laboratory - Centre for Plant health, North Saanich, BC, V8L 1H3 Canada; Summerland Research and Development Centre, Agriculture and Agri-Food Canada, Summerland, BC, V0H 1Z0, Canada; Translational Plant Biology, Department of Biology, Utrecht University, 3584CH, Utrecht, The Netherlands; State Key Laboratory of Integrated Management of Pest Insects and Rodents, Institute of Zoology, Chinese Academy of Sciences, Beijing 100101, China; Howard Hughes Medical Institute, University of California, Los Angeles, CA 90095, USA

**Keywords:** CRISPR, gene silencing, biotechnology, plant disease

## Abstract

Cassava production in sub-Saharan Africa is severely impacted by diseases. Most pathogens require interaction with host susceptibility factors to complete their life cycles and cause disease. Targeted DNA methylation is an epigenetic strategy to alter gene expression in plants and we previously reported that a zinc-finger fused to DMS3 could establish methylation at the promoter of *MeSWEET10a*, a bacterial susceptibility gene, and this resulted in decreased disease. Here, we attempt a similar strategy for cassava brown streak disease. This disease is caused by the ipomoviruses CBSV and UCBSV. These viruses belong to the family *Potyviridae*, which has been shown extensively to require host eIF4E-family proteins to infect plants and cause disease. We previously found that cassava plants with simultaneous knockout mutations in two *eIF4E* genes, *nCBP-1 and nCBP-2*, resulted in decreased susceptibility to CBSD. Here, we report successful simultaneous targeting of both promoters with methylation using a dCas9-DMRcd-SunTag system. However, in contrast to our previous work with *MeSWEET10a*, controls indicate that CRISPR interference is occurring in these lines and is sufficient for reduction of gene expression. Future research will use genetic crosses to segregate away the DNA methylation reagents and, if DNA methylation proves heritable, assess whether methylation alone is sufficient increase resistance to CBSD.

## Introduction

In plant-pathogen interactions, the manipulation of host processes by pathogens at the molecular level is critical for establishment of disease. These molecular interactions can involve interference with immune responses and co-opting or positively regulating host processes for the pathogen’s benefit. Host proteins that pathogens co-opt and use to promote infection are known as susceptibility factors. Some examples of well-studied genes encoding susceptibility (*S*) factors include *MLO* for powdery mildew (Büschges *et al*., 1997; Kusch and Panstruga, 2017), *SWEET* gene products for xanthomonads (Yang *et al*., 2006; Cohn *et al*., 2014; Mormile *et al*., 2024), and the *eIF4E*-family members for viruses of *Potyviridae* (Lellis *et al*., 2002; Duprat *et al*., 2002; Zaidi *et al*., 2018; Garcia-Ruiz *et al*., 2021). Disrupting pathogen interaction with susceptibility factors can confer disease resistance. This has previously been achieved through mutant *S* gene alleles or by using RNA interference (RNAi) to knockdown *S* gene expression (Zaidi *et al*., 2018; Garcia-Ruiz *et al*., 2021; Bastet *et al*., 2017; Takakura *et al*., 2018; Rubio *et al*., 2019).

The tropical root crop cassava is a staple for roughly 800 million people worldwide (Food and Agriculture Organization of the United Nations, 2018). Nearly half of cassava consumers reside in sub-Saharan Africa, where three major diseases, cassava bacterial blight (CBB), cassava mosaic disease (CMD) and cassava brown streak disease (CBSD) threaten cassava production (Bart and Taylor, 2017). CMD can be mitigated through the planting of resistant cultivars, but such cultivars are not widely available for CBB or CBSD (Fanou *et al*., 2018; Chikoti and Tembo, 2022). In cassava, *MeSWEET10*a and two pectate lyases are known *S* genes for CBB (Cohn *et al*., 2014; Cohn *et al*., 2016). DNA polymerase delta subunit 1 was recently described as a *S* gene for CMD (Wu *et al*., 2021; Lim *et al*., 2022) and CBSD requires interaction with two *eIF4E-*family genes, *nCBP-1* and *nCBP-2* (Gomez *et al*., 2019). We previously demonstrated that CRISPR-mediated mutagenesis at the *MeSWEET10a* promoter increased resistance to CBB (Elliott *et al*., 2024) and simultaneous knockout of *nCBP-1* and *nCBP-2* attenuated CBSD symptoms, including storage root necrosis (Gomez *et al*., 2019). For CBB a similar disease resistance outcome was achieved by directing DEFECTIVE IN MERISTEM SILENCING 3 (DMS3) to the *MeSWEET10a* promoter, resulting in *de novo* DNA methylation and suppression of pathogen-mediate gene induction (Veley *et al*., 2023).

CBSD is caused by independent or simultaneous infection by two potyvirids of the genus *Ipomovirus*, cassava brown streak virus (CBSV) and Ugandan cassava brown streak virus (UCBSV). The disease manifests as feathery vein chlorosis in leaves, brown streaking of stems, and corky necrosis that destroys the economically important storage roots (Tomlinson *et al*., 2018). While simultaneous knockout of *nCBP-1* and *nCBP-2* in the historical cultivar 60444 resulted in decreased disease compared to controls, these lines were not fully resistant. This lack of full resistance is possibly due to *eIF4E-*family redundancy, a hypothesis supported by protein-protein interaction data that demonstrates CBSV VPg can associate with the entire cassava complement of *eIF4E*-family encoded proteins (Gomez *et al*., 2019). Knocking out the entire *eIF4E-*family for CBSD resistance is infeasible due to their necessity for endogenous host protein translation. Knockdown, opposed to knockout, of susceptibility factors has been well documented to control potyviruses that exhibit *eIF4E-* or *eIF(iso)4E-*sub clade dependency (Bastet *et al*., 2017; Takakura *et al*., 2018; Rubio *et al*., 2019). However, the transferability of this method towards ipomovirus control is an open question, especially given potential differences in infection strategies used by potyviruses and ipomoviruses (Dombrovsky *et al*., 2014).

Our previous success with directing DNA methylation to the *MeSWEET10a* promoter inspired us to attempt epigenetic knockdown of members of the *eIF4E*-family genes in cassava. Here, we use a deactivated Cas9 protein to target the DOMAINS REARRANGED METHYLTRANSFERASE (DRM) catalytic domain to both *nCBP-1* and *nCBP-2* promoters for epigenome editing (Papikian *et al*., 2019). The resulting lines showed decreased gene expression. However, control lines lacking the DNA methylation reagent showed a comparable decrease in gene expression suggesting that dCas9 alone interferes with transcriptional machinery. This CRISPR interference (CRISPRi) effect has previously been reported to (Qi *et al*., 2013; Gilbert *et al*., 2014; Gardiner *et al*., 2022). Thus, the important next step will be to segregate away the DNA methylation reagents through genetic crosses and assess whether methylation is heritable and, if so, sufficient for increased resistance to CBSD.

## Results

### Production of transgenic cassava to knockdown susceptibility factor gene expression

We previously demonstrated that it is possible to direct methylation to a target region of the cassava genome (Veley *et al*., 2023). We hypothesized that targeting methylation to the *nCBP-1* and -*2* promoters would result in gene knockdown. To test this hypothesis, we used an epigenome editing tool comprised of three components, all present on a single T-DNA: (1) deactivated Cas9 with a C-terminally fused SunTag epitope (dCas9-SunTag), (2) *Nicotiana tabacum* DRM methyltransferase catalytic domain with its N-terminus fused to a single-chain variable fragment antibody that binds the SunTag epitope (scFv_SunTag_-DRMcd), and (3) a Pol II promoter driven tRNA-gRNA array (Figure 1a, c) (Čermák *et al*., 2017; Papikian *et al*., 2019). This tool has been used to re-establish heritable promoter methylation and subsequent repression of *FWA* at the *fwa-4* epiallele in *Arabidopsis*. Versions of this plasmid that omit either DRMcd (ΔDRMcd) or the gRNAs (ΔgRNA) were used as CRISPR interference (CRISPRi) and off-target methylation controls, respectively (Figure 1d, 1e). The CRISPRi construct was also evaluated for gene knockdown activity (Figure 1b).

**Figure 1.**
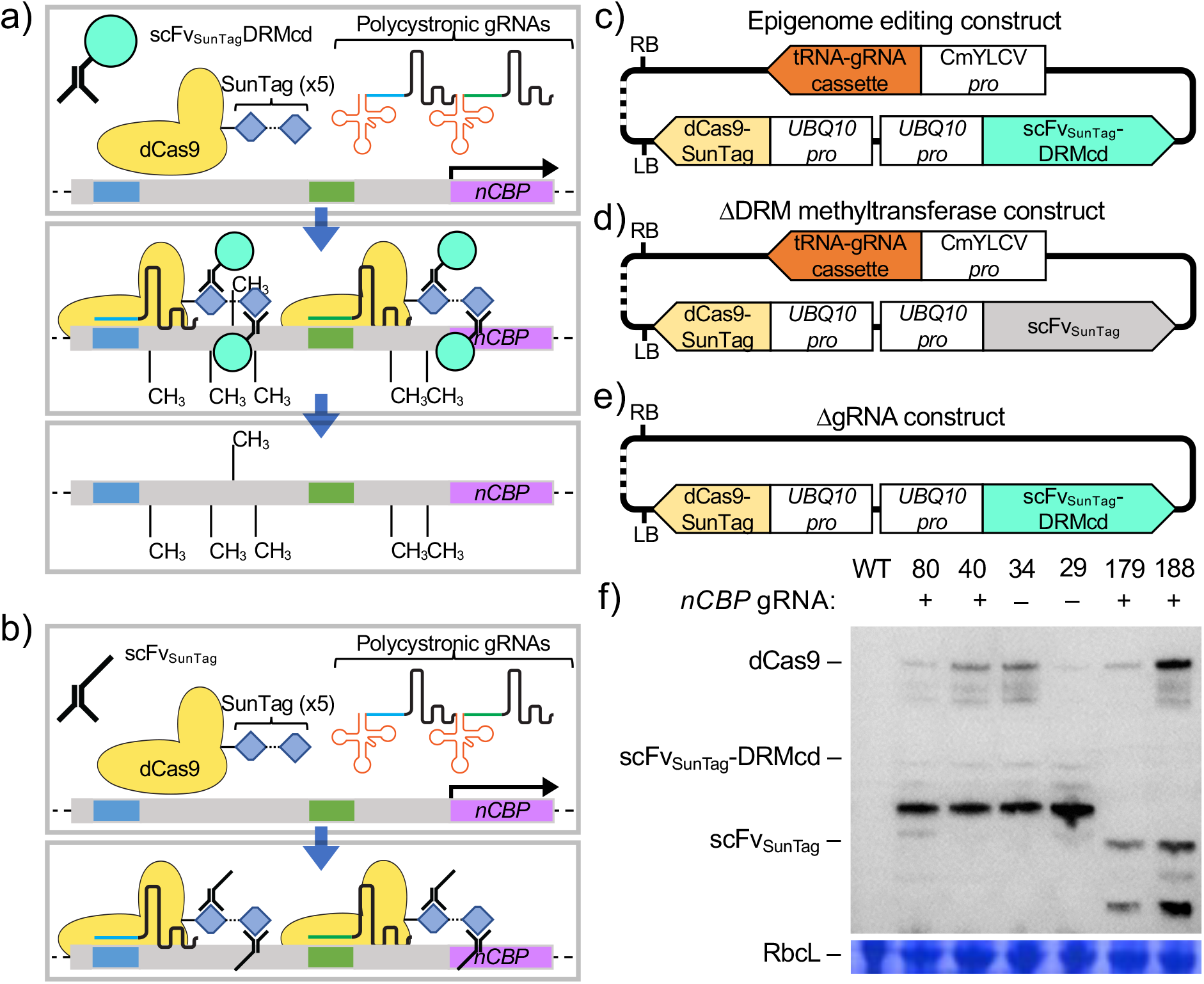
Strategy for dCas9-mediated gene knockdown a,b)Hypothesized mechanism of action for transcriptional knockdown via epigenome editing (a) or CRISPRi (b). c-e) Binary vector diagrams with color coded genes for dCas9-SunTag guided *de novo* methylation of DNA Fv_SunTag_-DRMcd. f)Transgene-encoded protein accumulation in all transgenic lines used in this study. HA-tagged dCas9-ag, scFv_SunTag_-DRMcd, and scFv_SunTag_ are all detected by western blot. Rubisco large subunit (RbcL) ted using coomassie brilliant blue staining of the same blot is presented as a loading control.

Two guide RNAs were used for each gene, *nCBP-1* and *nCBP-2*. Guide RNAs were designed to bind on either side of the putative first nucleotide of the respective 5’ UTR (based on Phytozome version 6) (shaded boxes in Figure 2a) (Bredeson *et al*., 2016). Following transformation into farmer-preferred cassava cultivar TME419, six relevant transgenic lines were recovered: ΔDRMcd lines #179 and #188; ΔgRNA lines #29 and #34; and lines containing the experimental epigenome-editing construct, #40 and #80. These lines were specifically selected for further study as the transgene-encoded proteins were expressed at similar levels (Figure 1f).

**Figure 2.**
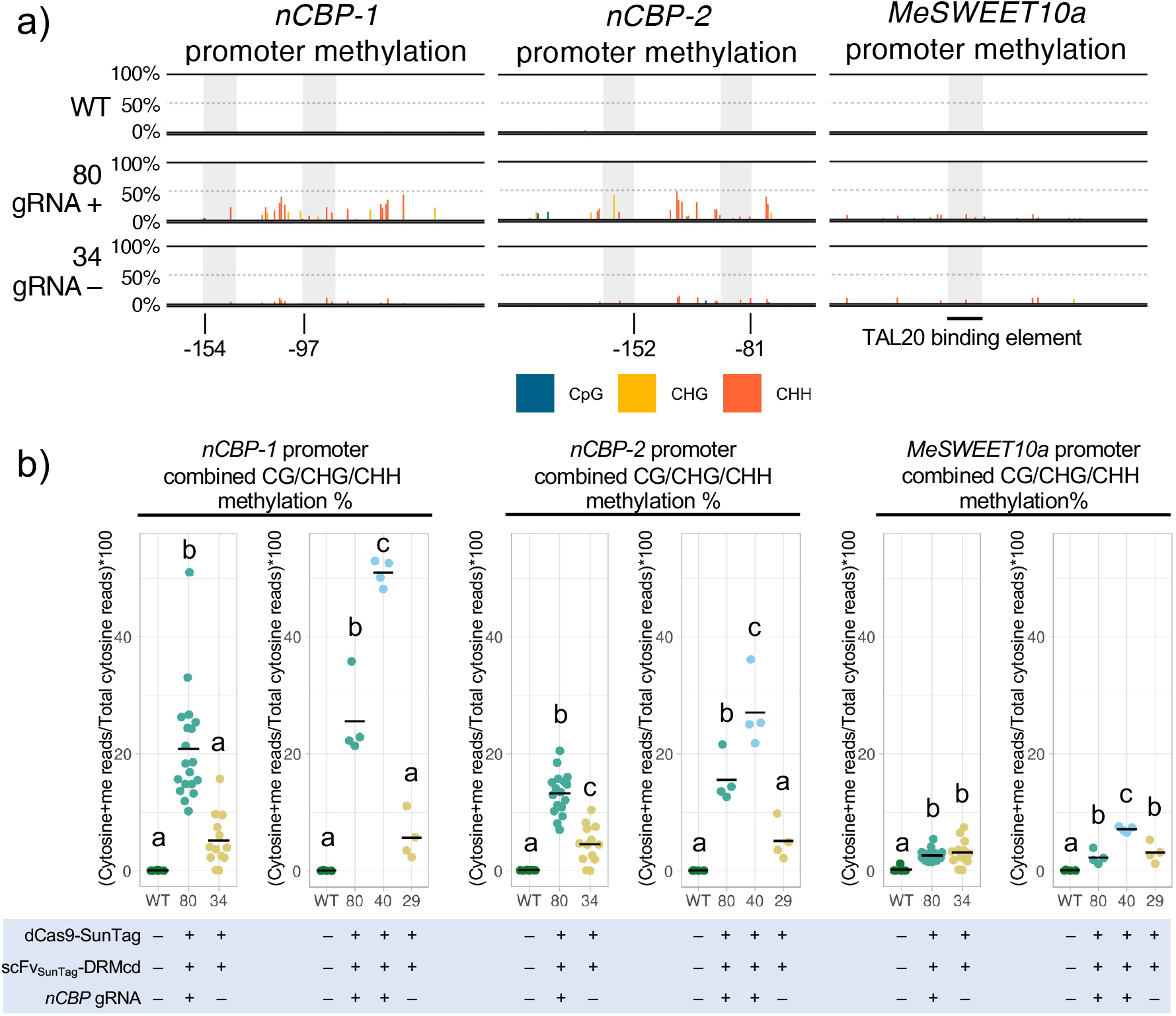
Targeting DRMcd to *nCBP* promoters produces modest levels of *de novo* CHH and CHG methylation a)Amplicon bisulfite sequencing of roughly 140-190 base pair windows in the *nCBP-1, nCBP-2*, and *SWEET10a* prmoters. These windows encompass the transcriptional start site and the right most nucleotide positions are -13, -69, and +1 base pair(s) relative to the first nucleotide of exon one, respectively. gRNA locations are shaded gray with the 5’ most nucleotide position relative to the start codon annotated. Line #80 and #34 both express dCas9-SunTag and scFv_SunTag_-DRMcd, but only #80 also expresses a gRNA cassette targeting both *nCBP-1* and *-2* b)Comparison of binned methylated-cytosine (cytosine+me) read percentage [100 x (methylated cytosine reads/total cytosine reads)] from within the BS-PCR seq window. Horizontal bar denotes mean methylation percentage. Statistical differences were detected by one-way ANOVA and post-hoc Tukey’s HSD, α=0.05.

### Targeted methylation to the *nCBP* promoters

Following identification of cassava transgenic lines carrying either the experimental or control vectors, we used amplicon-based bisulfite sequencing (ampBS-seq) to characterize the methylation patterns within the *nCBP* promoter regions. Wild-type TME419 plants exhibited no methylation within the window of interest while *de novo* methylation in the CHH and CHG contexts was observed in transgenic lines expressing dCas9-SunTag with the DRM catalytic domain (Figure 2a, b, S1, S2). Specifically, in lines #40 and #80, which contain gRNAs to target the dCas9-based machinery to the *nCBP* promoters, we observed significantly higher levels of cytosine methylation than in the control lines lacking gRNAs (lines #29 and #34) (Figure 2a, b, S1, S2).

To further confirm that the *de novo* methylation in lines #40 and #80 was specifically targeted to the *nCBP* regions, we characterized a third locus, the promoter of *MeSWEET10a*, a susceptibility factor for cassava bacterial blight disease. *MeSWEET10a* was chosen as we had previously characterized its promoter-methylation status using ampBS-seq (Veley *et al*., 2023). The *MeSWEET10a* promoter displayed weak levels of *de novo* methylation in all transgenic lines (Figure 2b, S3), indicating that while the presence of ectopic DRM catalytic domain is sufficient for low levels of non-specific *de novo* methylation, these levels are lower than when the enzyme is specifically targeted to a locus via gRNA-based recruitment of dCas9-SunTag.

### dCas9-SunTag occupancy of *nCBP-2* promoter results in transcriptional knockdown

As promoter methylation is usually associated with downregulation of gene expression, classically demonstrated at the *FWA* locus in *Arabidopsis* (Soppe *et al*., 2000; Bewick and Schmitz, 2017; Zhang *et al*., 2018), we investigated if targeting *de novo* methylation to the *nCBP-1* and *nCBP-2* promoters could downregulate these susceptibility genes. In an analysis of gene expression using RT-PCR, *nCBP-1* expression was not detected in wild type plants. Closer inspection of the *nCBP-1* gene model revealed that it was merged with the gene immediately downstream, Manes.09G140200 (Wilson *et al*., 2017; Gomez *et al*., 2019). Taking the correct, un-merged, gene model into account, *nCBP-1* expression is low or undetectable in all analyzed tissues, including cassava storage roots, stems, and leaves (Figure 3a). In tissues where *nCBP-1* transcripts were previously detected by RNAseq, *nCBP-2* is at least 30-fold more abundant. To confirm the expression of *nCBP-2* relative to *nCBP-1*, tissue culture plantlet leaves were analyzed by qPCR revealing that mean *nCBP-2* expression was 16-fold greater than that of *nCBP-1* (Figure 3b). Furthermore, 75% of *nCBP-1* cycle quantification values observed over multiple experiments are above 30, a consistent signature of low abundance transcripts that are difficult to quantify via qPCR (Taylor *et al*., 2019) (Figure S4). Given this result, we focused subsequent expression analyses on *nCBP-2* only.

**Figure 3.**
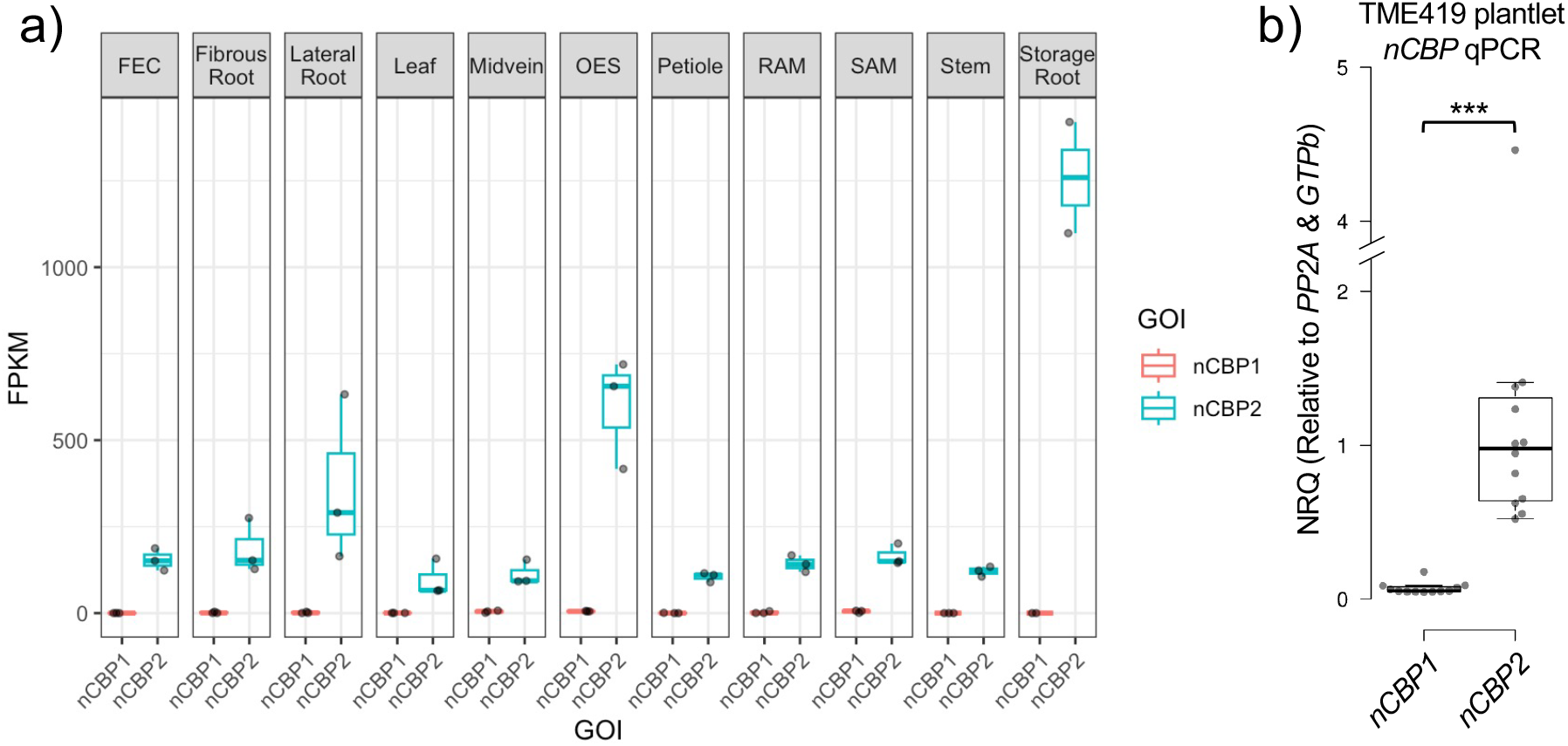
*nCBP-1* is weakly or not expressed in most tissue types *a)nCBP-1* and *nCBP-2* expression data extracted from the Bart Lab Cassava Atlas RNAseq browser. Gene expression levels are reported as fragments per kilobase of exon per million reads mapped (FPKM). b)qRT-PCR was used to detect *nCBP-1* and *nCBP-2* normalized relative quantities (NRQs), relative to *A* and *GTPb*, in plantlet leaves. The NRQs shown are relative to the geometric mean of *nCBP-2* measurements. Statistical differences were detected by Mann-Whitney U test, α=0.05, *** denotes p ≤ 0.001.

Expression of *nCBP-2* was significantly lower in epigenome edited lines #40 and #80 when compared to wild type and ΔgRNA lines (#29, #34) (Figure 4a, b). Therefore, *nCBP-*2 knock down is not due to off-target methylation. However, *nCBP-2* expression was also repressed in the CRISPRi control lines #179 and #188, which contain dCas9-SunTag, scFv_SunTag_, and the *nCBP-2* gRNAs but lack the DRM methylation domain, suggesting that *nCBP-2* knockdown in the epigenome edited lines may have been due to transcriptional interference (Figure 4c). Transcriptional repression due to steric hindrance by dCas9 of endogenous transcriptional machinery when targeted near transcriptional start sites has been previously observed in bacteria and human cells (Qi *et al*., 2013; Gilbert *et al*., 2014). Similarly, in plants, dCas9 has been fused to a transcriptional repressor domain for dCas9-based, targeted knockdown of gene expression (Pan *et al*., 2021). Both approaches are a form of CRISPR-interference (CRISPRi) (Pan *et al*., 2021; Qi *et al*., 2013). Our results indicate that gRNA-based targeting of dCas9-SunTag to a promoter region, even without fusion to a transcriptional repressor, is sufficient for significant knockdown of gene expression.

**Figure 4.**
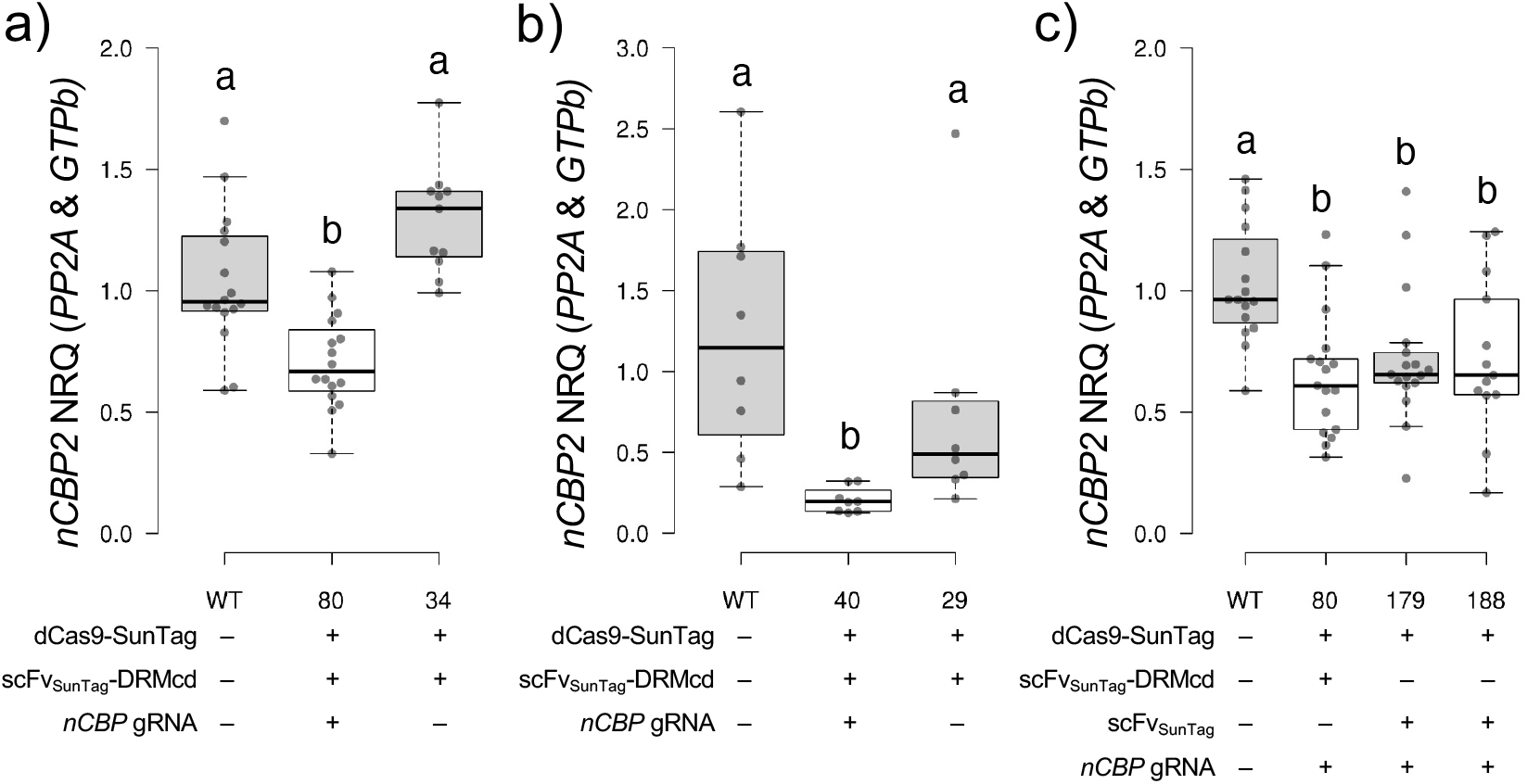
*nCBP-2* knockdown can be mediated by dCas9-SunTag without DRMcd *nCBP-2* expression levels in transgenic lines with and without gRNAs for targeted methylation (a, b) or with or without the DRMcd methylatransferase (c). qRT-PCR was used to detect *nCBP-2* normalized realtive quantities (NRQs) relative to *PP2A* and *GTPb*. The NRQs shown are relative to the geometric mean of wild-type measurements. The Kruskal-Wallis H test and post-hoc Dunn’s test using a Bonferroni corrected α were used in a-c) to detect significantly different mean ranks between paired ps.

## Discussion

We previously reported evidence that CBSV may recruit all cassava eIF4E-family proteins for disease. However, knocking out all eIF4E-family members simultaneously is not a viable strategy as these are critical host factors. We hypothesized that susceptibility factor knockdown may allow for the balancing of plant viability and disease severity. While susceptibility factor knockdown has been demonstrated to control disease caused by many potyviruses, this strategy had yet to be demonstrated for an ipomovirus with such broad eIF4E-family recruitment (Bastet *et al*., 2017; Takakura *et al*., 2018; Rubio *et al*., 2019). Here, we attempt to use targeted DNA methylation to knockdown expression of *nCBP-1* and *nCBP-*2. While we successfully achieved methylation at the target regions of the cassava genome, key control lines revealed that CRISPRi from the dCas9-SunTag fusion protein is sufficient to cause gene expression knockdown. Whether the level of *nCBP* promoter methylation achieved in this study is sufficient for heritable epigenetic knockdown will be further investigated by segregating away the epigenome editing tools.

An epigenome editing method is attractive as it may allow for transgene-free heritable knockdown of susceptibility factors. Thus far successful heritable epigenome editing has only been reliably accomplished at the *fwa-4* epiallele in *Arabidopsis thaliana* (Papikian *et al*., 2019; Ghoshal *et al*., 2021; Dubois and Roudier, 2021). While we were able to specifically enhance methylation at the *nCBP* promoters over wild type and ΔgRNA genetic backgrounds, the highest level of methylation observed at any cytosine for either *nCBP* promoter was 70% in the CHH context (Figure S1). Of the three possible methylation contexts, only CHH and CHG were observed. This is notable as CG methylation is thought to play a significant role in gene silencing and mediating inheritance of epigenetic marks (Ghoshal *et al*., 2021). The lack of CG methylation at the *nCBP* promoters in lines #40 and #80 is surprising given that our epigenome tool utilizes the catalytic domain of DRM, a *de novo* methyltransferase that methylates all three contexts (Papikian *et al*., 2019; Law and Jacobsen, 2010). In contrast, our previous work targeting DMS3, a non-catalytic component of RNA-directed *de novo* methylation machinery, to the *MeSWEET10a* promoter resulted in near 100% methylation of CG residues within the DNA motif required for *MeSWEET10a* upregulation by *Xanthomonas phaseoli* pv. *manihotis* (Veley *et al*., 2023). Promoter DNA methylation is not the only epigenetic mark associated with repression of gene expression; acetylation and methylation of certain lysine residues of histone 3 also control gene expression and have a complex interplay with DNA methylation (Du *et al*., 2015; Zhang *et al*., 2018). It is possible that existing histone epigenetic marks at the *nCBP* promoters antagonize DNA methylation while those at the *MeSWEET10a* promoter are permissive of DNA methylation.

The majority of CRISPRi studies in plants have focused on the development of dCas9-transcriptional repressor fusions (Piatek *et al*., 2015; Lowder *et al*., 2015; Vazquez-Vilar *et al*., 2016; Gardiner *et al*., 2022; Pan *et al*., 2021; Bhuyan *et al*., 2023; Wang *et al*., 2024). dCas9 alone has been shown to function as a transcriptional repressor in *Agrobacterium*-mediated transient assays in *Nicotiana benthamiana*, but has yet to be demonstrated in stable transgenic plants (Piatek *et al*., 2015; Vazquez-Vilar *et al*., 2016; Wang *et al*., 2024). Our approach differs from these transcription factor free fusions in that we use dCas9 fused to ten copies of the SunTag (GCN4) epitope that recruits ten copies of a single-chain variable fragment antibody fused to GFP (scFv_SunTag_). Furthermore, we simultaneously utilize two gRNAs in our target promoters. Our data clearly demonstrate that dCas9-SunTag/scFv_SunTag_ targeted to the transcriptional start site of *nCBP-2* can downregulate its expression (Figure 1b, 4c).

It remains to be seen whether *nCBP-2* knockdown is sufficient for attenuating CBSD symptoms. However, a knockdown strategy for CBSD resistance may provide some advantages compared to a traditional genome editing approach; the *eIF(iso)4E* and *nCBP* clades in cassava each consist of two homologs while *eIF4E* has just one. In *Arabidopsis, eIF4E* and *eIF(iso)4E* are individually dispensable for plant viability, but simultaneous mutations in both are lethal (Patrick *et al*., 2014). Assuming one intact *eIF4E* or *eIF(iso)4E* allele is required for viability, combining additional *eIF4E* and *eIF(iso)4E* mutations with those in the *nCBPs* for resistance requires several simultaneous CRISPR induced frameshift mutations all while preserving at least one *eIF4E*/*eIF(iso)4E* allele. Such a scenario may be difficult to obtain as insertion and deletion mutations induced by CRISPR are not always frameshifting, homozygous, or bi-allelic. Expanding our CRISPRi approach by targeting dCas9-SunTag or a dCas9-transcriptional repressor to additional *eIF4E*-family member promoters of interest would be an alternative method. For other purposes, CRISPRi and/or promoter methylation may prove advantageous compared to traditional RNAi-mediated knockdown in that dCas9 is targeted upstream of coding sequences, thus allowing for specific targeting of genes with close homologs sharing high sequence similarity. If CRISPRi were to be pursued further as an experimental or disease control strategy, future work might compare dCas9, dCas9-SunTag/scFv_SunTag_, and dCas9-repressor efficacy across multiple loci and also in the context of gRNA positional and dosage associated effects. As for targeted methylation, while this work clearly demonstrates feasibility of targeting specific loci with methylation, future work will need to understand heritability of *de novo* methylation and impact on gene expression in the absence of the DNA methylation reagents.

## Methods/Experimental Procedures

### Plant transformation and growth conditions

Transformation of cassava cultivar TME419 was performed as previously described (Veley *et al*., 2023). Transgenic lines were then maintained as *in vitro* plantlets (Taylor *et al*., 2012). For greenhouse growth, *in vitro* plantlets were moved to soil and allowed to establish on a misting bench before transfer to greenhouse conditions (Veley *et al*., 2023).

### Plasmid construction

gRNAs targeting the *nBCP* promoters (*nCBP-1*: CCGATGAATAAGAGCGCTAG, CGCTCTCAACTGTACTTCAT, *nCBP-2*: CACTCGATTGCAGATTTTTG, AGACGATGAAAGAGCCGAAG) were designed using CRISPOR (Concordet and Haeussler, 2018). gRNAs were assembled into pDIRECT_21C as described by (Čermák *et al*., 2017). The *CmYLCVpro::tRNA-gRNA* array was then amplified by PCR and cloned into the KpnI site of the dCas9-SunTag/scFv_SunTag_-DRMcd epigenome editing binary vector. The dCas9-SunTag/scFv_SunTag_ constructs were derived from dCas9-SunTag/scFv_SunTag_-DRMcd by utilizing BsiWI sites immediately flanking the *DRMcd* sequence.

### Methylation analysis

Cassava genomic DNA was extracted using the DNeasy Pro Mini kit (Qiagen), then prepared and processed for ampBS-seq as previously described (Veley *et al*., 2023). Primer sequences are provided in Table S1. Raw paired sequencing reads were merged and aligned to the following *nCBP* promoter sequences in CLC Genomics Workbench (Qiagen): *nCBP-1*, CCGATGAATAAGAGCGCTAGAGGTGAACGGTTTTCACCTGTCATCCTCTCTGGTTTCCGCTCTCAACTGTACTTCATAGGAAGCTATACTCGAATACAGAGACCTCCTATTAGAGCTTTTAGGATTTGGGGTTCAGCAAAG; *nCBP-2*, CCGAGTGCGGGAAATTTTATCACCGTTTCCAAACAGGCCACAAAAATCTGCAATCGAGTGA AATAAATGAAGCGTGTGTAGCTTTCCACTTCCTTTTCTCTCCGGTTTCCTCTTCGGCTCTTTCATCGTCTTCGCTGGACCT. Methylated cytosine reads were then called using the CLC Genomics Workbench epigenomic analysis plug-in.

### *nCBP* expression analysis

*In vitro* plantlet or greenhouse cassava leaf tissue was used for qPCR analysis of *nCBP-1* and *nCBP-2* expression. For *in vitro* plantlets, RNA was extracted using TRIzol (Invitrogen) in conjunction with the Direct-zol RNA Miniprep Kit (Zymo). For greenhouse grown plants, RNA was extracted using the Spectrum Plant Total RNA Kit (Sigma). cDNA was synthesized using the SuperScript III First-Strand Synthesis System (Invitrogen). qPCR was performed with SYBR Select Master Mix (Thermo Fisher) on CFX384 Touch (Bio-Rad) and QuantStudio 5 (Applied Biosystems) Real-Time PCR Systems. *nCBP* normalized relative quantities were calculated and analyzed as described in (Gomez *et al*., 2019). Primers used for qPCR are described in Table S1.

### Accession numbers

*nCBP-1*, Manes.09G140300; *nCBP-2*, Manes.08G145200; *GTPb*, Manes.09G086600; *PP2a*, Manes.09G039900; *MeSWEET10a*, Manes.06G123400

## Supporting information

Supplemental Table 1 - Primers used in this study

## Acknowledgements

We would like to thank the DDPSC greenhouse staff for their hard work and exceptional care of our plants. We are grateful to Nigel Taylor for graciously providing greenhouse space for our viral challenges. We would also like to thank Claire Albin for her extensive technical assistance and advice.

## Author Contributions

Conceptualization: J.C.C., S.E.J., R.S.B.; Methodology: Z.D.L., X.Z., S.F., B.G., J.G., M.W., J.C.C., S.E.J., R.S.B.; Investigation: Z.D.L., G.L.H., M.K.E., X.Z., S.F., B.G., J.G.; Visualization: Z.D.L., K.B.G., M.K.E.; Writing – original draft: Z.D.L.; Writing – review & editing: Z.D.L., G.L.H., M.K.E., X.Z., K.B.G., S.F., B.G., J.G., M.W., J.C.C., S.E.J., R.S.B.

