## Supplemental Figures 1-4 for "Targeting dCas9-SunTag to a susceptibility gene promoter is sufficient for CRISPR interference"

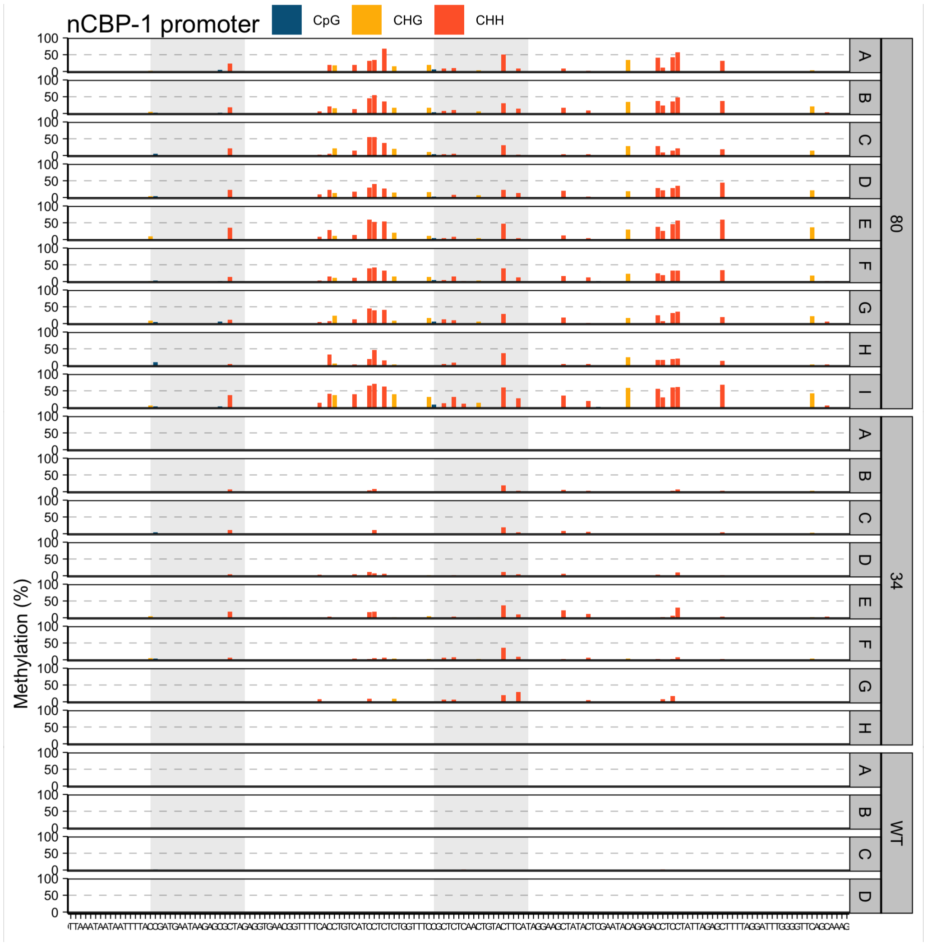


Figure S1. *nCBP-1* promoter cytosine methylation landscape

Percent methylated cytosines across the *nCBP-1* promoter ampBS-seq window for epigenome edited transgenic line #80 and the ΔgRNA control line #34. Each row within a genotype corresponds to an independent biological replicate.

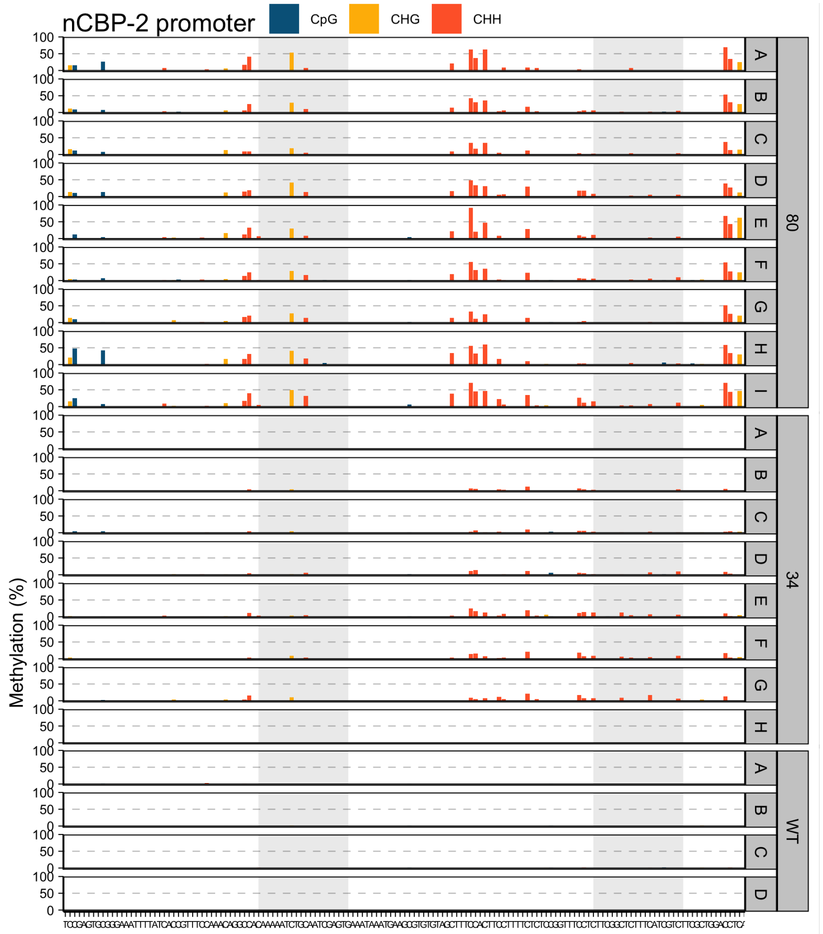


Figure S2. *nCBP-2* promoter cytosine methylation landscape

Percent methylated cytosines across the *nCBP-2* promoter ampBS-seq window for epigenome edited transgenic line #80 and the ∆gRNA control line #34. Each row within a genotype corresponds to an independent biological replicate.

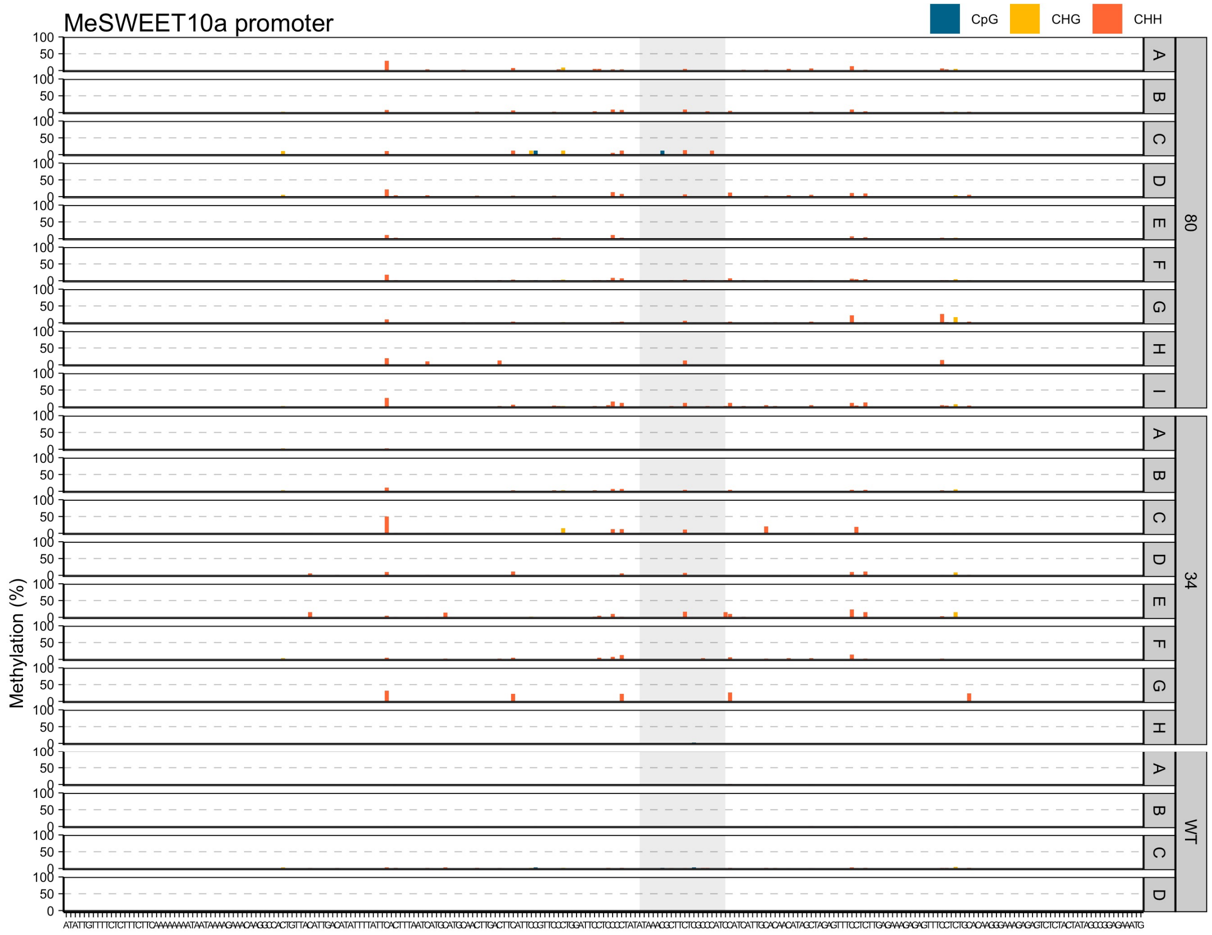


Figure S3. *MeSWEET10a* promoter cytosine methylation landscape

Percent methylated cytosines across the *MeSWEET10a* promoter ampBS-seq window for epigenome edited transgenic line #80 and the ∆gRNA control line #34. Each row within a genotype corresponds to an independent biological replicate.

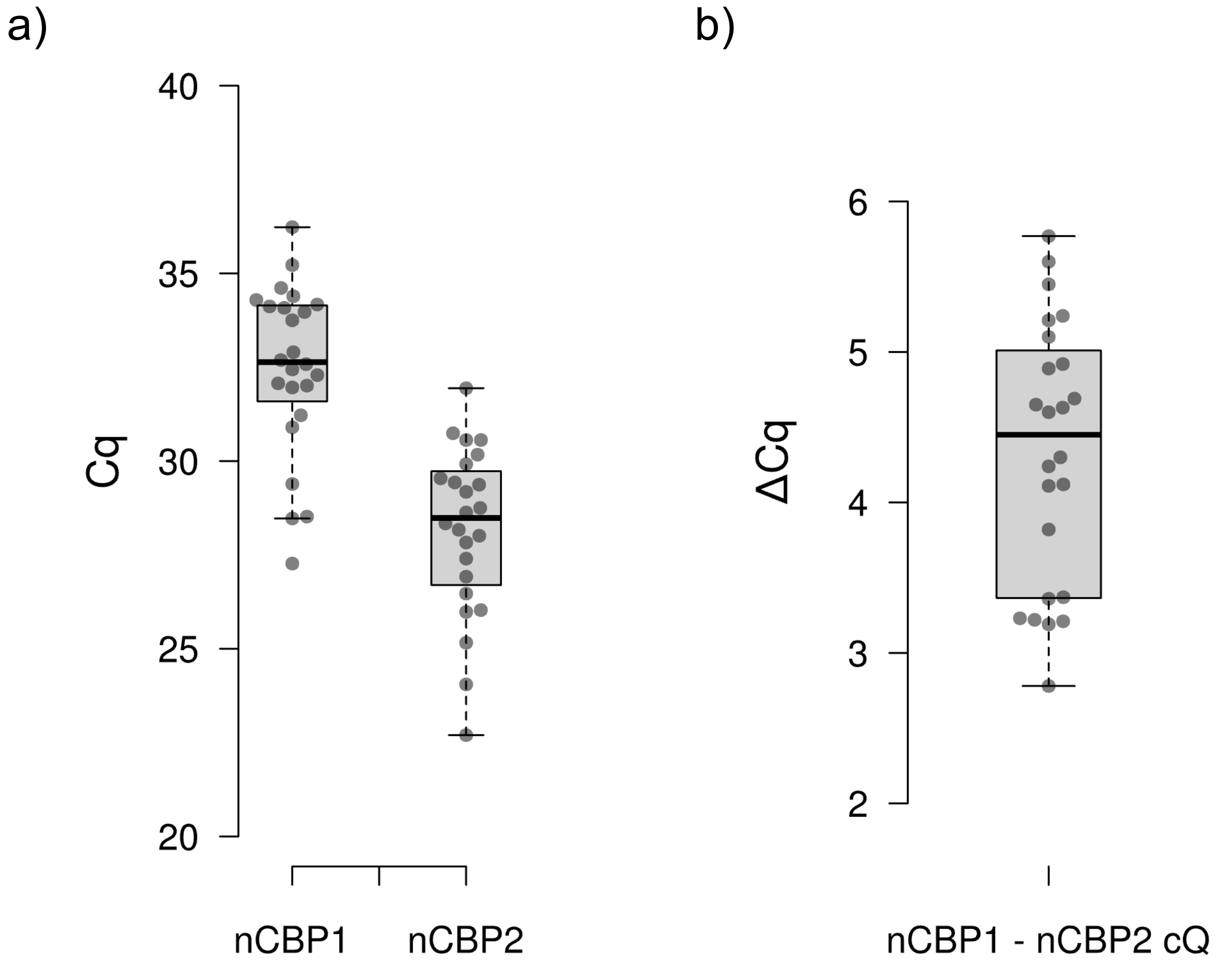


Figure S4.

a) Observed cycle quantitation (Cq) values for *nCBP1* and *nCBP2* across from across five qPCR experiments examining tissue culture leaf, greenhouse leaf, and CBSV infected storage root material. Threshold values for Cq selection were chosen based on those automatically selected by qPCR software for *nCBP-2* and then applied to *nCBP-1*.
b) Differences in *nCBP-1* and *nCBP-2* Cq from each tissue sample used in (a).
